# Noninvasive detection of bacterial biofilms using an insect olfactory brain-based gas sensor

**DOI:** 10.1101/2025.08.01.668178

**Authors:** Michael Parnas, Autumn K. McLane-Svoboda, Mariam Shahab, Camron Stout, Summer B. McLane-Svoboda, Elyssa Cox, Jonathan Hardy, Debajit Saha

## Abstract

Bacteria emit volatile organic compounds (VOCs) that can be targeted for disease detection. Biological olfactory systems have keen senses of smell, can detect VOCs at low concentrations, and are naturally adapted to classifying mixtures of VOCs as odors. Here, we employed locust (*Schistocerca americana*) olfactory neural circuitry to differentiate biofilm and planktonic cultures of *Pseudomonas aeruginosa* and *Staphylococcus aureus* using their odors. *In vivo* extracellular neural recordings were taken from the second-order olfactory processing center (antennal lobe) of locusts. The VOCs from biofilm cultures evoked distinct spiking responses compared to the planktonic cultures for both bacterial species. By analyzing the population neuronal responses, we classified individual bacterial biofilm vs. planktonic odors with up to 96% accuracy. The neural responses were highly discriminatory within the first couple of seconds of odor presentation and our analysis was conducted on less than five seconds of data, highlighting the potential of our biological sensor for real-time biofilm detection.

## Main

Biofilms can be 1000 times more resistant to antibiotics than planktonic bacteria^1^ and account for 80% of bacterial infections in humans^2^, including wound, implant, and lung infection. These multicellular, heterogenous colonies form on solid surfaces and create a complex extracellular environment, constituting, but not limited to, a polysaccharide matrix filled with extracellular DNA (eDNA)^3^. The combination of the extracellular matrix and heterogenous nature makes it extremely difficult for antibiotics to completely eliminate a biofilm. Some almost dormant bacteria buried within the matrix, known as persisters, have slower metabolisms that enable them to survive and recreate the biofilm post-treatment^4^. Without proper and timely treatment, biofilms are the perfect safe havens for bacteria to survive, evolve, and maintain chronic infections.

Current methods for detection and diagnosis of bacteria mostly rely on culturing blood or other biological samples to grow a bacterial culture *in vitro*. These tests take a few days to grow the sample in a lab for results and require skilled personnel^5^. While faster detection methods such as qPCR and ELISA have been developed, these do not distinguish between biofilm and planktonic, and they can only test a single targeted disease at a time and^6–8^. Matrix-assisted laser desorption–time of flight (MALDI-TOF) mass spectrometry has been used to identify specific bacterial strains^9^ and can differentiate between biofilm and planktonic cultures^10,11^. However, MALDI-TOF requires the isolation of bacterial species for culturing, which could result in invasive patient sampling and longer analysis times or a very high infection load to directly analyze patient samples such as blood and urine^12^. Volatile detection techniques offer a non-invasive and rapid alternative to traditional diagnostics by analyzing the gas-phase volatile organic compound (VOC) concentrations emitted by a sample, such as breath, to determine the presence of a disease. Gas chromatography-mass spectrometry (GC-MS), the gold standard for VOC analysis, and other analytical chemistry techniques have been used to detect and differentiate biofilm^13–17^ and planktonic^18–29^ bacterial species based on their VOC profiles. Unfortunately, these analytical techniques require sample preparation, trained personnel to operate, and are too time-consuming for rapid or real-time diagnosis^30^. While analytical instruments are always improving, their main strength is sensitive individual compound identification^31^. To overcome these hurdles, electronic noses (e-noses) use biological olfactory principles such as cross-selective chemical sensors and pattern matching analysis with combinatorial coding^32,33^. E-noses have been used to detect bacteria *in vitro*^34–38^, *in vivo*^39–42^, and even distinguish between biofilm-forming and planktonic bacteria. Though e-noses are portable, easy to use, and provide rapid VOC analysis, each device is designed to detect a few specific VOC mixtures associated with a target disease and lacks the specificity and sensitivity of biological olfaction^33,43–45^.

Biological olfaction, one of the oldest evolved senses, has solved the problem of detecting VOCs rapidly, with broad sensitivity, and reliably across both time and changing environments^46,47^. Our review offers an extensive dive into the existing literature of biological olfaction for disease detection through behavior, component-based sensors, and whole-brain sensors^48^. Dogs are well known for their keen sense of smell, whether for security purposes^49,50^, or medically relevant detection of diseases^51,52^ such as cancers^53,54^ and bacterial infections^55,56^. Invertebrates also have an extremely sensitive sense of smell, on par with dogs, and *Caenorhabditis elegans* has been used in behavioral assays to detect VOCs associated with bacteria^57–59^. Unfortunately, behavioral assays from these species have binary outputs (e.g., presence or absence), which limit their usefulness when multiple diseases could be present. Our previous work has tapped into the neural responses within locust and honeybee brains to detect endometriosis^60^, and oral^61^ and lung^62,63^ cancer cell cultures, without the limitations of behavioral assays. Rather than mimicking biological olfaction or relying on behavior, here we are directly using the locust (*Schistocerca americana*) olfactory system to detect biofilm and planktonic cultures of *Pseudomonas aeruginosa* (PA) and *Staphylococcus aureus* (SA) from their VOCs.

## Results

### Bacterial biofilm odors were classified using a biological olfactory brain

We used the locust olfactory system to differentiate between biofilm and planktonic bacterial cultures from two different species. The bacteria were cultured in customized air-tight T25 flasks (Methods: Bacteria culturing) to produce five different odor flasks: *P. aeruginosa* biofilm (PAbio), *P. aeruginosa* planktonic (PAplank), *S. aureus* biofilm (SAbio), *S. aureus* planktonic (SAplank), and Luria-Bertani (LB) broth as a control (**Fig. 1a**). Meanwhile, a locust was prepared for extracellular electrophysiology recordings from the antennal lobe (AL) (Methods: Electrophysiology, **Fig. 1b**). After all preparations were completed, the headspace from the odor flasks were presented to the locust antennae in a pseudorandom order using an olfactometer (Methods: Odor stimulation, **Fig. 1c, Extended Data Fig. 1**). We used a pseudorandom order to eliminate temporal biases, such as the possibility of a stronger response to the first odor presented than the last odor. The locust antenna, analogous to vertebrate noses, interact with VOCs within the surrounding gaseous environment. Sensilla, tiny (about 10μm long) hair-like follicles that cover the exterior of the antennae, have pores for VOCs to enter the antennal lymph^64^. There, VOCs can interact with odorant receptors (ORs) on the membranes of olfactory receptor neurons (ORNs) to initiate an electrical signal down the antennal nerve and into the AL. Individual ORNs express one of the over 100 ORs, but each OR can be sensitive to multiple VOCs^65^. Locusts, and most biological olfactory systems including vertebrates, combine the different response profiles of many ORs to enable a drastic increase in the number of unique odors they can detect. As an example, locusts using combinatorial coding with a binary, on/off, paradigm for each OR, could theoretically encode over 1.2 × 10^30^ (2^n^ – 1) unique odors from just 100 OR^66^. Once the neural signal reaches the AL, it is processed by a dense interconnected network of excitatory projection neurons (PNs) and inhibitory local neurons. The PNs then output to higher-order brain centers (e.g., Mushroom Body) for learning and decision making. Interestingly, there are over 100,000 ORNs within each antennae but only ~800 PNs within an AL, connecting with over 300,000 neurons in the mushroom bodies^46,67^. Due to this bottleneck, all the olfactory information is carried by only 800 PNs, which is why we target the PNs for data collection. We collected voltage traces, the raw data, that contain responses from multiple PNs using an electrode carefully placed in the AL (**Fig. 1b,c**). The voltage traces were spike sorted (Methods: Spike Sorting) to isolate the response pattern of each PN. The PN responses were then combined for analysis and high-dimensional ‘fingerprints’ were created for each of the odors based on the population neural response (**Fig. 1d**). Using relatively simple pattern matching and Euclidean distance metrics we then quantified the classification performance of the locust olfactory system for the tested odors (**Fig. 1e**).

**Figure 1:**
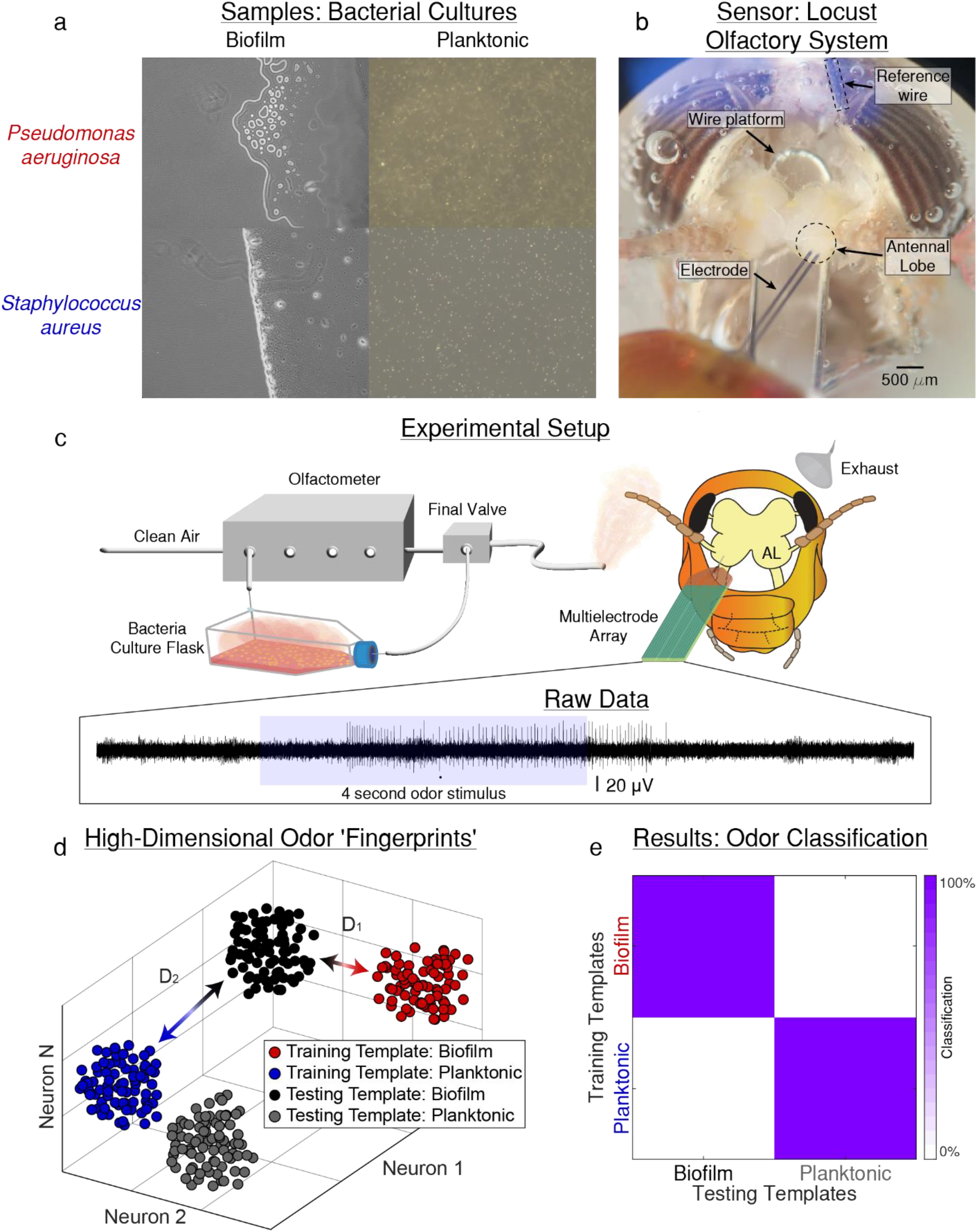
Schematic of the experiment delivering bacterial odors to a locust olfactory sensor and subsequent data analysis for classification. **(a)** *Pseudomonas aeruginosa* (PA) and *Staphylococcus aureus* (SA) were cultured as biofilms (left) and planktonic (right) to prepare the four odor samples. All cultures were prepared in Luria-Bertani (LB) broth which was used as the fifth odor for control. **(b)** Locust, *Schistocerca americanus*, were used as a biological sensor to detect odors. Locust antennae (analogous to vertebrate noses) detect volatiles in the air. They were prepared via surgery which removed the exoskeleton and exposed the AL while maintaining the antennal nerve connections to the antennae. A wire platform was placed under the brain to support it and prevent motion during data recording. A reference wire was placed within the headspace but not touching the brain. The electrode was carefully placed within the AL using a micromanipulator. **(c)** Headspace from the bacteria culture flasks were delivered to the locust antennae using an olfactometer for precise odor delivery timing (**Extended Data Fig. 1**). A cone pulling a slight vacuum was placed behind the locust to quickly clear the odors after delivery. Raw data was collected as voltage recordings using the electrode and time matched to the odor delivery timing. **(d)** All of the recorded neurons were combined to give the entire population of neurons responses to each odor. Each neuron’s response was analyzed as an orthogonal dimension which together yields a high-dimensional response ‘fingerprint’ for each odor. The data was split between training and testing templates for each odor, and then the testing templates were assigned to whichever training template minimized the high-dimensional Euclidean distance. **(e)** The results of the high-dimensional classification were summarized in a confusion matrix indicating the percentage of testing templates assigned to each of the training templates.

### Neural responses can differentiate between biofilm and planktonic cultures of *Pseudomonas aeruginosa*

The initial investigation focused on the responses of single PNs within the AL to bacterial odors. This was to determine whether the locust olfactory system could detect these bacteria. The raw voltage traces acquired from extracellular electrophysiology recordings are the amalgamation of neural responses from a small region of the brain, which can include multiple neurons or specifically, for the brain region we recorded from, multiple PNs (**Fig. 2a**). In this representative voltage trace from a single recording location, there are spikes caused by neuron action potentials. These spikes could be attributed to a single neuron or to many different neurons. However, the voltage spike shapes of individual neurons are unique, based on proximity to the electrode and neuron morphology^68^, which can be leveraged to spike sort, e.g., assign each of the voltage spikes to individual neurons. Using a spike sorting algorithm^69^ we can create models for each of the spike shapes within a voltage trace and apply these models to all of the voltage traces obtained during that recording. The output is the spike times assigned to individual PNs (32 PNs identified). To visualize a single PN’s responses, the spike times for each of the five trials of odor presentation can be plotted as a raster, and the trials can be averaged together into a peri-stimulus time histogram (PSTH) (**Fig. 2b**). For this specific neuron, notice that prior to the odor presentation (blue window) there was very little spiking that then increased after odor presentation started and rapidly returned to the pre-stimulus baseline after odor presentation stopped. This PN had a slightly larger steady-state (~2-4 seconds after stimulus onset) and off (immediately after stimulus offset) response to the biofilm than the planktonic odor and both of these responses were much larger than the response to the LB control. These differences are important for classification analyses.

**Figure 2:**
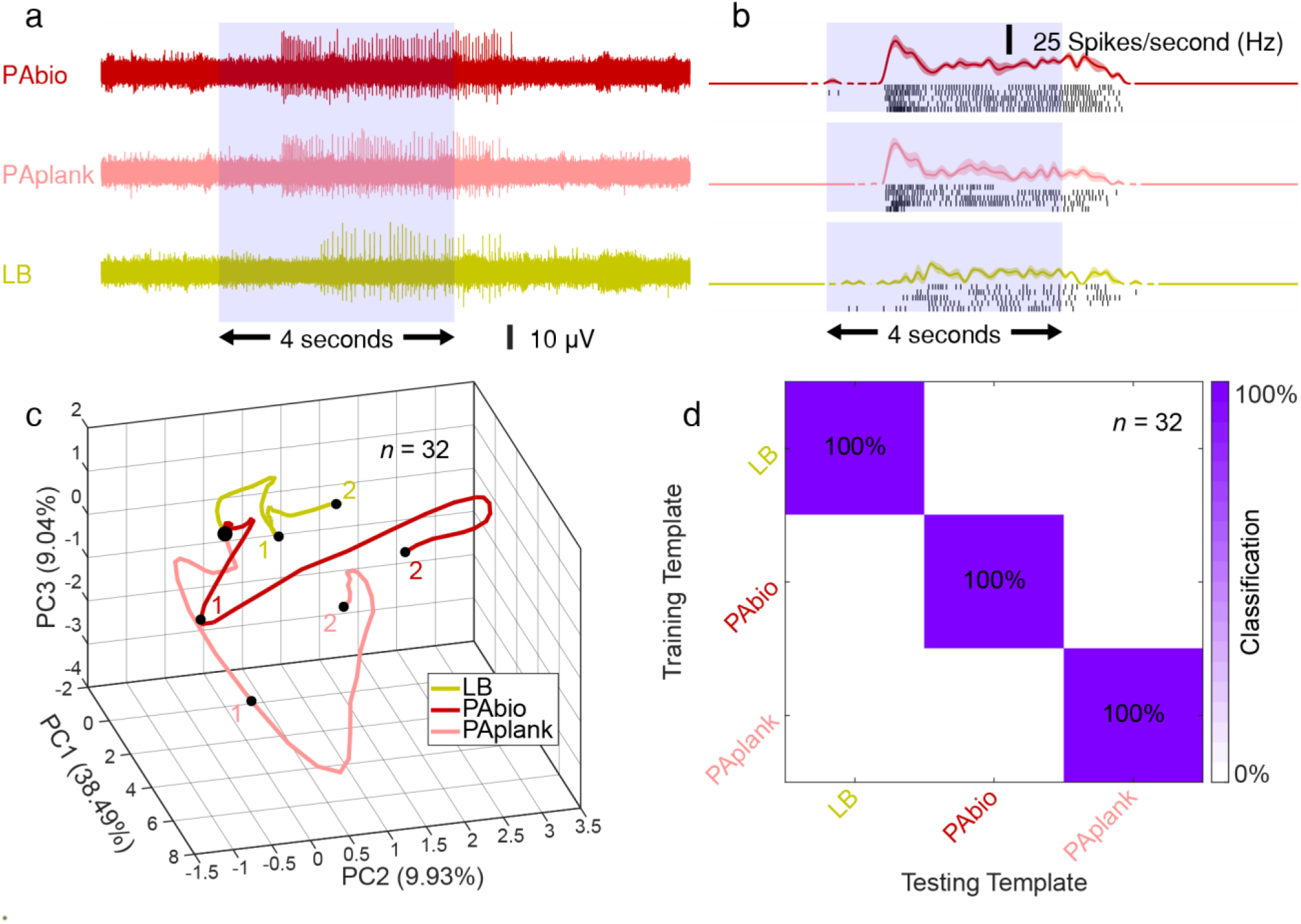
Neural responses can differentiate between biofilm and planktonic cultures of *Pseudomonas aeruginosa*. **(a)** We show representative voltage traces of extracellular neural responses to the bacterial odors from the AL. There are three odors, biofilm (PAbio, dark red), planktonic (PAplank, light red), and Luria-Bertani broth (LB, yellow). The light blue box corresponds with the four-second odor presentation window. Spikes within the voltage trace are due to PN activity. **(b)** The recording location shown in **panel a** was spike sorted to identify individual PN spike times from the voltage trace. Here, we show a single PN’s odor-evoked responses with raster and peri-stimulus time histograms (PSTHs). Each black line in the raster plot indicates a single spiking event from the PN, and each row of black lines indicates a single trial (five trials total). The PSTH is the trial-averaged response with the S.E.M. indicated by the shaded region. The light blue box corresponds with the four-second odor presentation window. **(c)** Population PN trajectory plots created using PCA dimensionality reduction to reduce from 32 dimensions (one dimension for each neuron) to three for visualization. The trajectories trace the evolution of the neural response across the first two seconds of the stimulus starting at the origin. The small numbers within the odor space denote seconds after the onset of odor delivery. There is visual differentiation between the population neural responses of each odor within these three dimensions. **(d)** Population PN responses are classified using a leave-one-trial-out (LOTO) cross-validation with no dimensionality reduction and summarized in confusion matrices. The spiking responses across the four second stimulus duration are separated into 50ms time bins (80 total). For each odor, one trial is used for testing while the remaining four are averaged together as the training data. This creates a testing template and a training template for each of the five odors at each of the 80 time bins. Within each time bin, the testing templates are compared to the training templates and assigned to the template that minimizes Euclidean distance. In a winner-take-all (WTA) approach, each testing trial is assigned a single classification using the mode of the time bin classifications. The training templates are on the y-axis and the testing templates are on the x-axis. Here, we can see that all trials were correctly classified, yielding 100% accuracy for the detection of biofilm vs. planktonic *P. aeruginosa*.

To examine the entire population of 32 identified PNs, we conducted principal component analysis (PCA) to visualize the population neural trajectories (**Fig. 2c**). PCA, an unsupervised learning algorithm, was used to reduce 32-dimensional PN responses to the three principal components (PCs) that explain the largest variance in the dataset. PN spike times are binned into 50ms non-overlapping time bins. This matches the 20Hz oscillations found within the locust AL which is thought to allow for information integration prior to projection to higher-order brain centers^46^. This 50ms binned data is used in all subsequent analyses. The time-binned data of all 32 PNs are transformed using PCA, and then the data points are connected in sequential order to show the temporal evolution of the neural response. To better understand the relationship between the temporal neural responses and the PCA neural trajectories, we checked the responses from two neurons to two different odors (**Extended Data Fig. 2a**). These spike timings were plotted with the neurons on separate axes (**Extended Data Fig. 2b**). Using just two neurons, we can visualize the relationship between temporal odor-evoked responses without using dimensionality reduction. This visualization can then be extended out to all 32 neurons using PCA discussed above (**Extended Data Fig. 2c**).

Trajectories that travel further from the origin (e.g., PAbio and PAplank) indicate larger neural responses with higher spiking rates and differences in the trajectory angles indicate different sets of neurons are responding^70^. Within most olfactory responses there is a transient phase associated with odor onset, which is the most discriminatory, followed by a steady state that is still different from the baseline but not as discriminatory as the transient phase. Here, we focused the analysis on the transient response during the first two seconds of odor stimulus. We observed clear differences in the trajectories for all the odors, revealing that the population PN responses are unique for each bacterial odor. This PCA transformation explains over 50% of the variance in the dataset (obtained by summing PC1, 2, and 3) and therefore is used for qualitative purposes only.

We used a leave-one-trial-out (LOTO) analysis to quantitatively assess the classification accuracy of the PN responses using previously described methods^61,62^. As there are three odors, this analysis is a three-way test in which each testing template is compared to all three training templates and assigned using minimum Euclidean distance; completely random classification should yield 33% accuracy. Spikes were binned into non-overlapping 50ms time bins and then classified using time-matched training and testing templates. In a winner-take-all (WTA) approach, the mode of time bin classifications was used to classify each trial, and achieved 100% accuracy (**Fig. 2d**). The WTA approach avoids the stochastic trial to trial variation within the 50ms bins that can be observed in the raster plots (**Fig. 2b**). Using neural information from only 32 PNs, we were able to discriminate between biofilm and planktonic cultures of *P. aeruginosa*.

### *Staphylococcus aureus* biofilm and planktonic cultures were also differentiated

A key strength of biological olfaction is its sensitivity to a broad swath of odors. The locust sensor, with no genetic or chemical manipulation, not only responded to *P. aeruginosa* but also *S. aureus* odors. There was an odor-evoked response at the single PN level to *S. aureus* biofilm and planktonic odors as seen in a representative neuron (**Fig. 3a,b**). At the population neural response level, SAbio and SAplank project in different directions within the neural response space indicating distinct responses (**Fig. 3c**). And, using a LOTO analysis, we were able to differentiate between the neural responses to these odors (**Fig. 3d**). Only one of the five trials, 20%, of SAplank was misclassified to SAbio. The misclassification between LB and both *S. aureus* odors could be due to some of the odor-evoked response being from the control.

**Figure 3:**
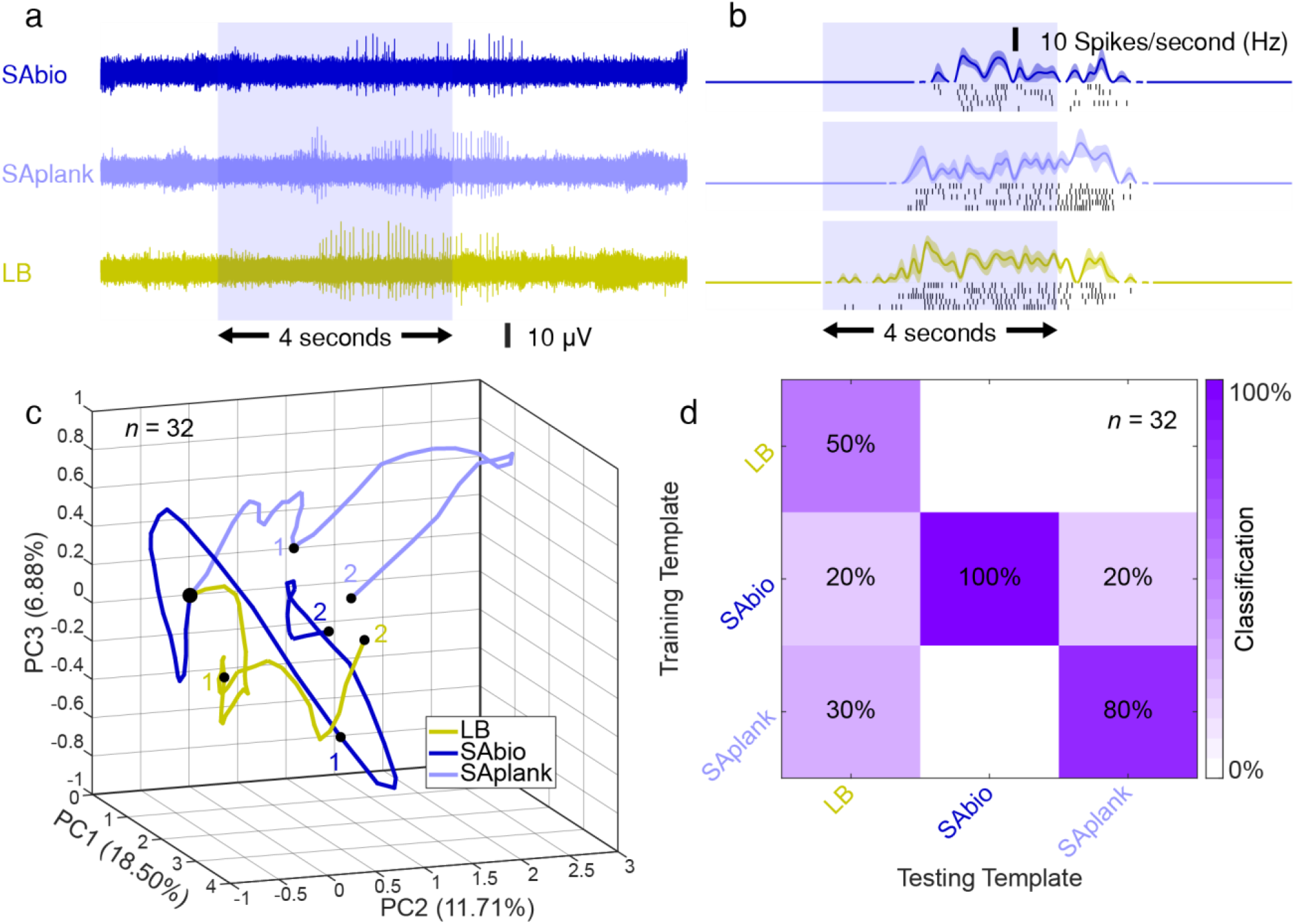
The locust olfactory system also responds to odors from *Staphylococcus aureus*. **(a)** Representative voltage traces of extracellular neural responses to the bacterial odors from the AL. There are three odors, biofilm (SAbio, dark blue), planktonic (SAplank, light blue), and Luria-Bertani broth (LB, yellow). **(b)** The recording location shown in **panel a** was spike sorted, and a single PN’s odor-evoked responses were visualized using raster and peri-stimulus time histograms (PSTHs). **(c)** Population PN trajectory plots created using PCA dimensionality reduction across the first two seconds of the stimuli. The small numbers within the odor space denote seconds after the onset of odor delivery. There is visual differentiation between the population neural responses of each odor within these three dimensions. **(d)** Population PN responses are classified with LOTO cross-validation using a high-dimensional Euclidean distance metric. Here, we can see that all the trials were correctly classified, yielding 100% accuracy for the detection of biofilm vs. planktonic *P. aeruginosa*.

### Locust projection neurons can encode the odors of biofilm cultures from two different bacterial species, *Pseudomonas aeruginosa* and *Staphylococcus aureus*

We combined the neural responses to both bacterial species into a single analytical test to classify biofilms. PCA of the transient odor response, first two seconds of odor stimulus, shows distinct trajectories for each of the five odors with the *P. aeruginosa* odors (red) projecting towards the top and *S. aureus* odors (blue) projecting towards the bottom (**Fig. 4a**). Over 50% of the variance is explained in the first three PCs. A quantitative classification using LOTO achieves 88% accuracy of the entire four-second odor stimulus (**Fig. 4b**). PAbio and SAbio were both classified with 100% accuracy. One trial (20%) of SAplank was misclassified as SAbio.

**Figure 4:**
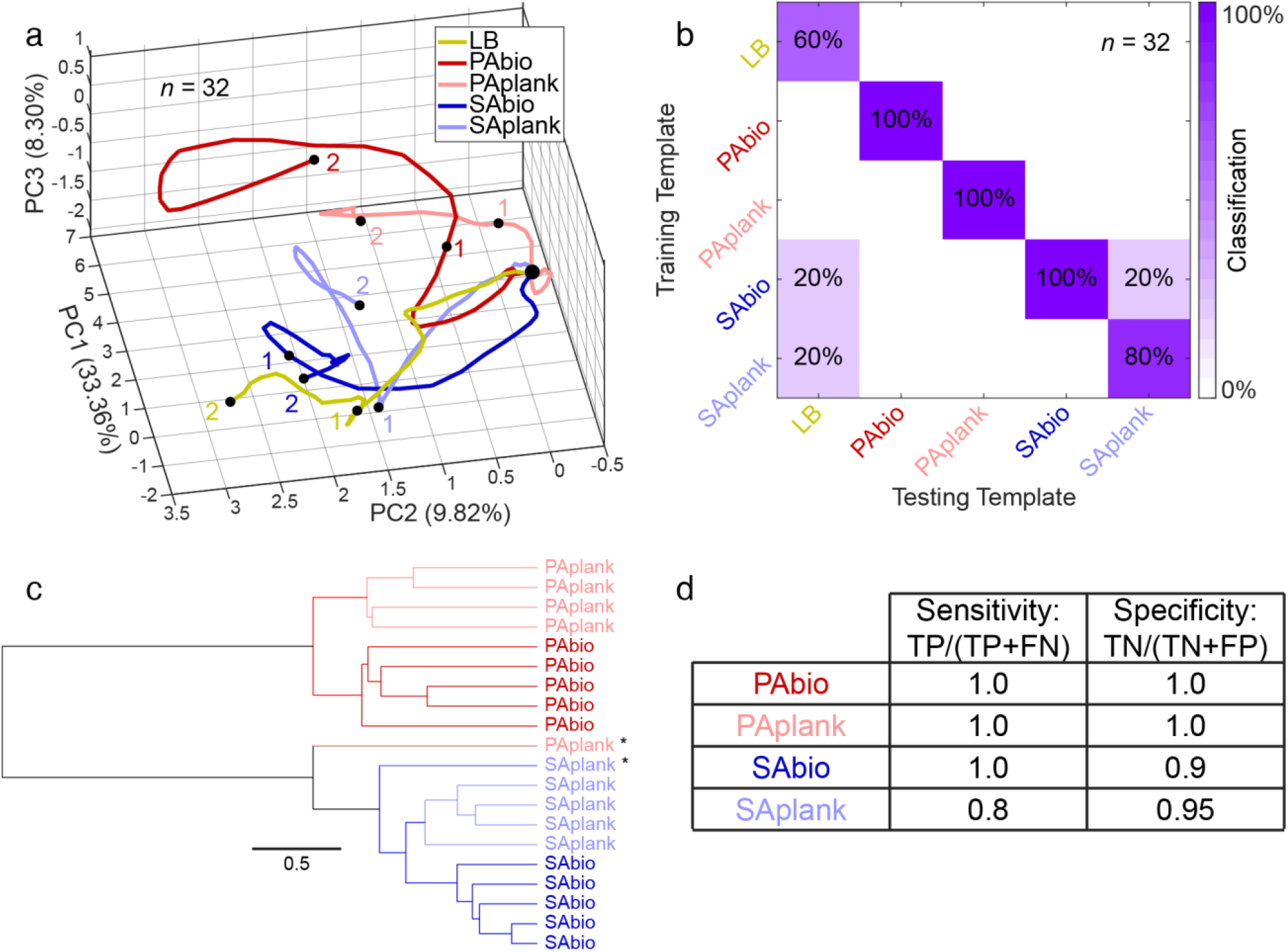
Population neural responses uniquely encode each of the bacterial odors. **(a)** The population neural responses in PCA space all project in unique trajectories across the first two seconds of stimuli. The large differences in the trajectory angles indicate different neural subpopulations encode for the different bacterial species (red: *Pseudomonas aeruginosa* and blue: *Staphylococcus aureus*). The biofilm (dark) and planktonic (light) odors of each species also elicit distinct trajectories from each other. The small numbers within the odor space denote seconds after the onset of odor delivery. **(b)** Using the Euclidean distance, LOTO analysis across the four second odor presentation window, the *P. aeruginosa* (dark red) and *S. aureus* (dark blue) biofilms were both classified with 100% accuracy. *P. aeruginosa* planktonic (light red) was also classified with 100% accuracy. One out of five trials (80%) of *S. aureus* planktonic (light blue) was misclassified to *S. aureus* biofilm. The control, LB broth (yellow), was confused with the *S. aureus* odors in two out of five trials. **(c)** A high-dimensional hierarchical clustering using Ward’s method shows that the largest difference in odors is between bacterial species with the next major split between biofilm and planktonic odors of the same bacterial species. This highlights that differentiating between bacterial species is easier, but locusts can still do the much more difficult task of differentiating biofilm from planktonic bacterial cultures, and they can do this for both bacterial species. **(d)** A one-vs-rest analysis for specificity was implemented in which each odor is its own correct classification, and all other odors are incorrect, thereby creating multiple binary scenarios for calculating sensitivity and specificity. We see very high sensitivity and specificity for each of the odors.

To further understand the neural encoding of bacterial odors, we performed a hierarchical clustering analysis (**Fig. 4c**). We compared the time-matched 50ms bins across the entire four second stimulus duration using Euclidean distance in 32 dimensions. We see the largest difference in the data is between bacterial species, unsurprisingly, with a sub-splitting of the data between biofilm and planktonic cultures. These results suggest that the locust VOC sensor can more easily differentiate between bacterial species, however, it can still detect the more subtle differences between biofilm and planktonic cultures.

The sensitivity, calculated as the ratio of true positive test results (TP) over the presence of the odor (TP + FN), and specificity, the ratio of true negative test results (TN) over the absence of the odor (TN + FP), for each of the individual bacterial odors was high (**Fig. 4d**). PAbio, PAplank, and SAbio all achieved 1.0 sensitivity and both Pabio and PAplank had 1.0 specificity. The locust sensor reliably detected each of the bacterial odors.

### The spatiotemporal classification occurs rapidly and robustly after odor delivery

We then tested the classification ability using different subsets of the data to investigate the transient response period. We applied the WTA analysis to different time window sizes: 0.25s (5 time bins), 0.5s (10 time bins), and 1s (20 time bins) (**Fig. 5a**). As expected, the classification accuracy is low during the pre-stimulus time period, indicating there are no biases. However, after odor-onset during the transient response, the accuracy rapidly increases to 82-91% before slightly decreasing during the steady state response. These results are robust across different time windows and mirror the high overall accuracy of 88% achieved using the entire 4s window (**Fig. 4b**). The highest classification accuracies all center on ~1s after stimulus onset (**Fig. 5b**).

**Figure 5:**
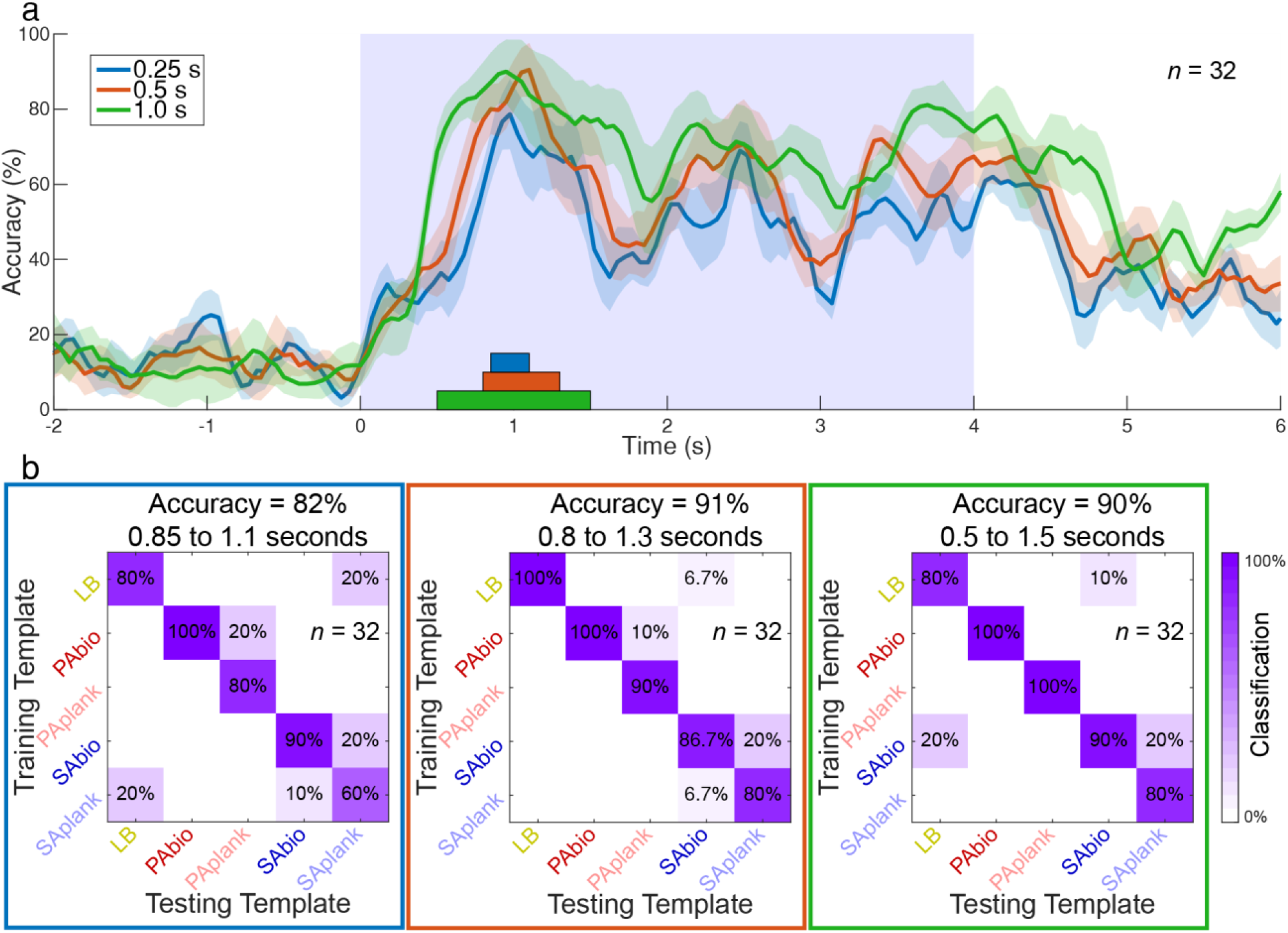
Accurate classification occurs rapidly after odor onset and with less than one second of data. **(a)** The LOTO accuracies for different time windows are plotted. The S.E.M. of the accuracies across the five trials is the shaded region. Using a time range of 0.25s (blue), or only five 50ms time bins, we have roughly 20% accuracy during the pre-stimulus windows (−2s to 0s), which we expect for a completely random 5-way classification test. However, very rapidly after odor presentation the accuracy climbs to over 80% after about a second. The same trends hold for the 0.5s (red) and 1.0s (green) time ranges. The colored rectangles indicate the time windows with the highest accuracy for each time range. **(b)** The highest accuracy confusion matrices for each of the time ranges (0.25, 0.5, and 1s). The highest accuracy time windows for the three time ranges are centered around 1s after stimulus onset, which aligns with the transient neural response. As more data is included, the accuracy increases from 82% to over 90%; slightly better than including neural response data from the entire four-second stimulus (**Fig. 4b**).

### Modeling the capabilities of the locust olfactory system for biofilm detection

In any classification model, thresholds between positive and negative classifications need to be properly and conscientiously chosen. Previously, we used the minimum Euclidean distance to classify odors; this effectively places the threshold exactly in the middle between any two odors (**Extended Data Fig. 4**). Here, we varied this threshold by biasing the model towards or away from the PAbio (**Fig. 6a**) and SAbio (**Fig. 6b**) odors. At each threshold, the sensitivity and specificity can be calculated to create a receiver operating characteristic (ROC) curve. We also sampled random subsets (10, 20, or 30 PN) of the population of 32 recorded PNs and saw that the sensor performance trended upwards with the inclusion of more neurons for both biofilm odors. We modeled this trend as an exponential by comparing the area under the curve (AUC) with the number of neurons used in the model (**Fig. 6c,d**). We assumed that zero neurons would yield a completely random binary classifier and that as the number of neurons approaches infinite, the classification should approach perfection. The exponential regression models fit the data with R^2^ values over 0.95. Finally, to determine the best classification thresholds to use for model generation, we calculated the Youden’s Index (YI), a combination of sensitivity and specificity (**Fig. 6e,f**). The maximum YI, or best classification performance, for both biofilm odors occurs at a threshold very close to 1, which is the same as the minimum Euclidean distance. Therefore, using the minimum Euclidean distance in future odor classification problems would be a good approximation for the optimal classification threshold.

**Figure 6:**
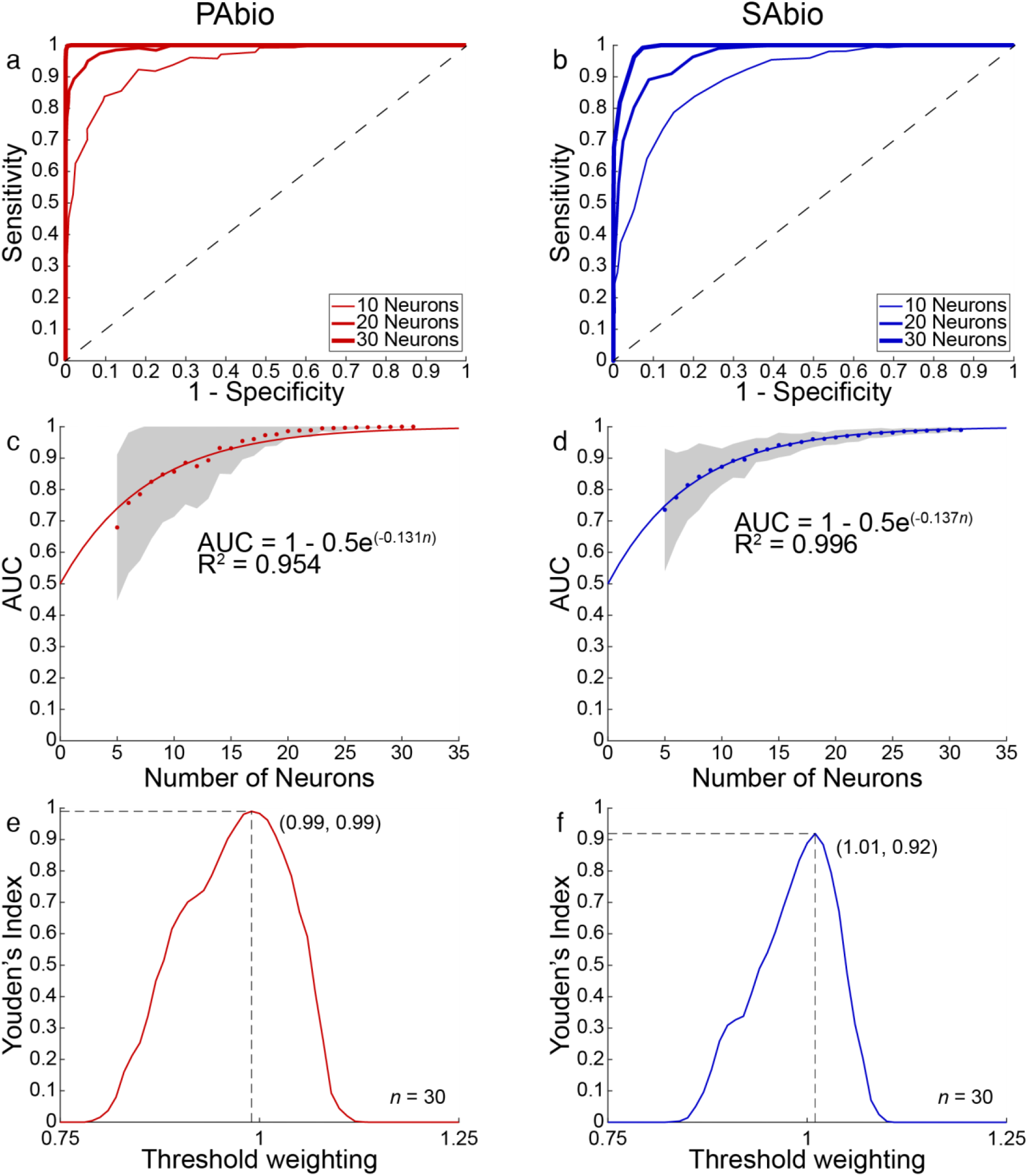
Modeling the locust neural olfactory sensor’s capabilities. **(a)** Receiver operating characteristic (ROC) curve of the locust sensor’s detection of *P. aeruginosa* biofilm produced by varying the classification threshold (Methods: ROC analysis, **Extended Data Fig. 4**). Out of 32 total recorded PNs, random subsets of neurons (10, 20, and 30) were sampled and a clear trend of better sensor performance correlates with more neurons. For each number of neurons, 100 random samples were chosen and averaged together to produce the curves. The dashed line indicates the performance of a completely random binary classifier. **(b)** For *S. aureus* biofilm, there is a similar trend of better sensor performance as the number of neurons increase. **(c-d)** To further explore the trends seen in **Fig. 6a,b**, the area under the curve (AUC) was compared with the number of neurons sampled. Each dot is the average AUC for 100 random subsamples of the 32 recorded neurons, and the grey shaded region is one standard deviation. An exponential curve was fitted to the data, assuming that the x-intercept is 0.5, the classification is completely random when there is no neural data, and that the curve should approach an AUC of 1, or perfect classification, as you increase the number of neurons. From both these models, an AUC of over 0.9 is achieved with just 15 neurons. The coefficients of determination (R^2^) were both greater than 0.95, indicating a strong fit to the model. **(e-f)** Youden’s Index, a linear combination of sensitivity and specificity, is calculated at different values of the threshold weighting. A threshold weighting of 1 is the same as minimum Euclidean distance. A threshold weighting greater than 1 makes unknown odors less likely to be classified as the odor of interest, PAbio or SAbio. This decreases sensitivity. However, a threshold weighting less than 1 makes unknown odors more likely to be classified as the odor of interest, which decreases specificity instead. For both PAbio and SAbio, the highest Youden’s Index occurs very close to a threshold of 1, or minimum Euclidean distance, and is over 0.9.

## Discussion

This work uses the neural responses from the olfactory brain centers of locusts to differentiate between odors produced by biofilm and planktonic cultures of PA and SA. VOCs generated through normal cellular metabolism are excreted into the LB and then enter the gaseous headspace. While there are shared metabolic mechanisms between species of bacteria, there are clear differences in the metabolomes^71,72^. These differences result in changes to the VOC profile or odor which are detectable via analytical chemistry techniques^13–29^, e-noses^34–42^, and biological olfactory systems such as in this work. Changes in gene expression, the proteome, and the metabolome are also evident between biofilms and planktonic bacteria^73–77^. To our knowledge there has been no study comparing the similarity of volatilomes between bacterial species to the similarity between biofilm and planktonic states. However, we observed that the biological olfactory system sensed the biofilm vs. planktonic states more similarly, while still clearly differentiating them, than the two bacterial species, *P. aeruginosa* vs. *S. aureus*.

Analytical chemistry techniques identify individual compounds. However, the identification of VOCs of interest is based on which compounds give the largest signal, which may not be the most important compounds for disease identification. E-noses require fore-knowledge of the VOCs of interest for sensor material selection, which is generally based on the data produced by the analytical chemistry techniques before using pattern matching for discriminating between disease and healthy states. On the other hand, our biological olfactory sensor does not require prior understanding of compounds associated with different diseases, instead, odors are presented to a completely naïve locust sensor, and patterns within the neural responses are matched to classify diseases. We harnessed the incredibly broadly tuned yet still sensitive insect antenna and gathered the neural output from the first-order odor encoding center of the brain, the AL. The neural outputs provided unique spatiotemporal patterns associated with each of the different bacterial culture headspaces. Then, using a relatively simple pattern matching approach, we could identify and classify each of the entire complex VOC mixtures as one of the bacterial odors. This approach has been previously used to detect different cancers and is generalizable to many different odors^60–63^.

A limitation of this study is the LOTO analysis, which does not use independent training and testing data. However, we did a separate classification analysis with non-overlapping training and testing trials that yielded similar results to the LOTO analysis. While biofilms are inherently more resistant to antibiotics than planktonic bacteria, and here we differentiated between those, different strains of these bacteria have already developed resistance to some antibiotics^78^. We expect that trying to classify two different strains of the same bacterial species (e.g., methicillin-sensitive vs. methicillin-resistant SA) will be more difficult, but this has been done by e-noses, which provides confidence that it can also be done by the locust sensor^34^. Here, we only collected neural recordings from 32 PNs, roughly 4% of the total PNs in the locust AL, and still achieved high levels of classification. Gathering information from a larger fraction of the PNs should yield more sensitive classifications. Finally, we did not explore the impact of comorbidities such as cystic fibrosis on our detection capabilities. *P. aeruginosa* is a common persistent infection associated with cystic fibrosis patients, in part due to biofilm formation^79^. We conducted all tests on *in vitro* bacterial cultures to achieve better control and differentiate biofilm vs. planktonic; in the future we will use breath studies with more variability and the presence of comorbidities.

Since the early 20^th^ century antibiotics have extended the average human lifespan by over two decades and completely changed modern medicine, however, there is now an incredible lack of novel antibiotics being produced^80^. Also, current antibiotics are overprescribed, similar to opioids in the 1990s, as preventative measures in many cases without a proper diagnosis of a bacterial infection, much less the type of bacterial infection^81,82^. In combination, the lack of novel drugs and overprescription of current drugs, is leading to potential microbial epidemics^83^. A rapid, point-of-care, and non-invasive VOC detection device that could inform a doctor’s choice by identifying the presence of biofilms and differentiating bacterial species would alleviate the overprescription or incorrect prescription of antibiotics.

## Methods

### 1. Electrophysiology

*In vivo* extracellular neural recordings were conducted on post-fifth instar, but pre-mating stage, locusts (*Schistocerca americana*) of either sex raised in a crowded colony.

#### Surgery

Surgery to access the antennal lobes (AL) for neural recordings followed previously published methods^84^. First, locusts were immobilized by amputating the legs, sealing the amputation sites with a tissue adhesive (Vetbond, 3M), and securing the locust’s thorax to a custom surgical bed using electrical tape. Antennae were immobilized by threading them through small polyethylene tubes (Intramedic Microbore Tubing, 427420) within a poly-vinyl chloride 18 AWG wire insulation jacket that were then secured to batik wax pillars. A batik wax bowl was then built connecting the pillars to the head, starting above the mandibles and encompassing the compound eyes, with the rim extending just above the top of the head. Prior to making any incision, the wax bowl is filled with physiologically balanced locust saline solution at room temperature^85^. The exoskeleton and glandular tissue covering the brain was removed. The gut, which the brain rests on, was also removed to prevent brain movement and then the brain was supported with a batik wax-coated wire platform connected to the batik wax bowl (**Fig. 1b**). Finally, the thin membrane sheath covering the AL was removed following a short treatment with protease.

#### Recording

A commercial Neuronexus 16-channel silicon probe (A2×2-tet-3mm-150-150-121) was electroplated to achieve impedances between 200 to 300kΩ. The electrode was inserted about 100μm into the AL. A silver chloride ground wire was placed within the head of the locust but not touching the brain to complete the circuit through the saline solution. While recording, a small saline drip (10 mL/hr) was used to prevent loss of saline solution in the wax bowl due to evaporation. During data collection, voltages were sampled at 20kHz and digitized by an Intan pre-amplifier board (C3334 RHD 16-channel headstage). These digital signals were transferred to the Intan recording controller (C3100 RHD USB interface board) and then stored on a PC using the Intan GUI. Six locusts were used for electrophysiology experiments.

### 2. Odor stimulation

Odors were delivered to the insect antennae via a commercial olfactometer (Aurora Scientific, 220A) (**Fig. 1a, Extended Data Fig. 1**). During trials, air flow through the three mass flow controllers, MFC1, MFC2, and MFC3, were held constant at 120, 80, and 200 sccm, respectively. Total flow to the insect antennae was also held constant by controlling the final valve to deliver the 200 sccm from MFC3 or the combined 200 sccm from MFC1 and 2. The constant air flow to the insect antennae mitigates potential confounding neural responses from mechanosensory neurons. At the start of a trial, the final valve is configured to deliver clean air from MFC3. Five seconds prior to odor delivery, the final valve continues to deliver air from MFC3 while valves 1 and 2 open, and valve 3 closes, to divert the 80 sccm of air from MFC2 through the bacterial culture flask and to prime the airlines up to the final valve. During odor delivery the final valve switches and delivers the combined airflow from MFC1 and 2 to the insect antennae for four seconds. After four seconds, the final valve reverts to the original position, delivering clean air from MFC3, valves 1 and 2 close, and valve 3 opens to reduce VOC loss from the bacterial culture headspace. A 6” diameter funnel pulling a slight vacuum is always behind the locust to clear the VOCs after odor delivery. All airlines use PTFE to reduce VOC interactions. Clean air is provided by a compressed air cylinder at 12 PSI. During each experiment, a replicate of the same odor panel was used: *Pseudomonas aeruginosa* biofilm (PAbio), *Pseudomonas aeruginosa* planktonic (PAplank), *Staphylococcus aureus* biofilm (SAbio), *Staphylococcus aureus* planktonic (SAplank), and lysogeny broth (LB). The order of odor delivery was pseudorandomized, and each odor was delivered five consecutive times with a 1-minute inter-stimulus interval.

### 3. Bacteria culturing

For the preparation of biofilms, overnight cultures of Xen5 (*Pseudomonas aeruginosa*) and Xen36 (*Staphylococcus aureus*) were grown in Luria-Bertani (LB) broth. The cultures were then diluted 1:100 in fresh LB broth, and 5 mL was seeded into 25 cm^2^ customized tissue culture flasks. Prior to bacteria seeding, the flasks were altered by inserting inlet and outlet 20-gauge needles through the cap and vertically through one of the upper corners opposite the cap. Both needles were then secured using a low-volatile, two-part epoxy (Permatex, 84101), which also sealed the holes around the needles. The epoxy was allowed to cure for longer than 24 hours. At all times, except during odor delivery, the inlet and outlet needles were capped to prevent VOC loss. Because *Pseudomonas* forms biofilms at the air-liquid interface, it needs an edge upon which to attach, so the flasks were tilted at 30°. The flasks were then placed in a 37°C incubator overnight. To verify that biofilms had formed, bioluminescent images were obtained with the IVIS instrument (Perkin-Elmer Inc.). For planktonic cultures, 400 mL of overnight culture in LB broth was added to 20 mL of LB broth and grown in a 250 mL Erlenmyer flask at 37°C shaking at 200 RPM for 1.5 hours. Optical density was taken, and 10^7^ colony-forming units were added to 1 mL in a tissue culture flask with a needle port in the cap. Immediately before experiments were performed, the planktonic bacteria were removed from the biofilms, and 1 mL of LB broth was added to prevent drying. All flasks contained 1 mL of LB broth, including the LB broth control.

### 4. Spike sorting

Intan neural data was imported into MATLAB for high-pass filtering using a Butterworth filter to remove components below 300Hz. Data was then spike sorted in Igor Pro using previously described methods^69^. Each tetrode from the Neuronexus electrode was considered an independent recording and spike sorted. The detection threshold to identify spiking events was maintained between 2.5-3.5 standard deviations (SD) of baseline fluctuations. Single PNs were incorporated into the dataset if they passed three criteria: 1) cluster separation > 5 SD, 2) inter-spike intervals < 10%, and 3) spike waveform variance < 10%. A total of 32 PNs from six locusts passed all the criteria.

### 5. Hierarchical clustering

Spikes were binned into 50 ms non-overlapping time bins. A baseline spike rate for each neuron was calculated as the average spike rate during the two seconds preceding the odor stimulus and then subtracted from the dataset. Then the Euclidean distance in high-dimensional space (32 neurons yield 32 dimensions) between observations (5 trials x 5 odors = 25 observations) was calculated for each time point during the stimulus presentation (4s time window/50 ms time bins = 80 time points). The distances across time were averaged for each pair of observations to yield 300 distance values (25 observations choose 2 = 300). Linkages were calculated using Ward’s method to produce a dendrogram (**Fig. 4c**). Hierarchical clustering was done using custom written MATLAB (R2020b) code.

### 6. Dimensionality reduction analyses

We performed principal component analysis (PCA) to reduce the dimensionality of the dataset for visualization purposes^61,62^. Spikes were binned into 50ms non-overlapping time bins. A baseline spike rate for each neuron was calculated as the average spike rate during the two seconds preceding the odor stimulus and then subtracted from the dataset. We then averaged the five trials together. Each of the neurons is a separate dimension (**Extended Data Fig. 2**), so we performed PCA dimensionality reduction on the combination of all five odors (LB, PAplank, PAbio, SAplank, and SAbio) to reduce from 32 dimensions to the three principal components (PCs) that explained the largest variance (PC1, 2, and 3). Each of the odors was then plotted along these three PCs with the data connected in temporal order to create the neural trajectories (**Fig. 4a**). The trajectories were smoothed using a third-order IIR Butterworth filter (Half Power Frequency = 0.15) and shifted to begin at the origin. PCA was done using custom written MATLAB (R2020b) code.

### 7. Quantitative classification analysis

To assess the bacterial odor classification performance, we performed a *leave-one-trial-out* (LOTO) cross-validation analysis^61,62^. Spikes were binned into 50ms non-overlapping time bins. A baseline spike rate for each neuron was calculated as the average spike rate during the two seconds preceding the odor stimulus and then subtracted from the dataset. Four out of the five trials were averaged together to yield high-dimensional, temporal, training neural data (32 neurons x 80 time bins) for each of the odors (**Fig. 4b**). The left-out trials from each odor were used as testing data. Therefore, for each of the 50ms time bins, there were five points in high-dimensional space for training and another five points for testing; one training and one testing point for each odor. The testing points were compared with all of the training points and assigned to the odor that minimized the Euclidean distance (**Fig. 1d**). Then, we iterated through each of the trials, each time leaving a different one out for testing and using the average of the remaining four as training data. So, for a four-second analysis window, there are 80 time bins x 5 odors x 5 trials = 2000 time bins that we are attempting to classify. Additionally, a winner-take-all approach was implemented by using the results of the bin-wise classification to vote. Here, the most common classification assignment, or the mode, of each trial was used to assign the entire testing trial to one of the training odors (**Fig. 1e**).

We also used completely separate training and testing datasets by averaging two random trials together to create the training templates and using the remaining three trials as three separate testing templates (**Extended Data Fig. 3a**). We assessed the robustness of these results by conducting this analysis on all possible combinations of two training trials with the accuracy results summarized in a histogram (**Extended Data Fig. 3b**).

To investigate the impact of different analysis time windows the prior winner-take-all approach was applied across different time windows of the dataset (**Fig. 5**). Here, three different time window sizes, 0.25s (5 time bins), 0.5s (10 time bins), and 1s (20 time bins), were tested across a continuous moving window. The averages of the trial-wise classification accuracies were plotted with the S.E.M. indicated by the shaded region (**Fig. 5a**). Each line indicates the accuracy at the center of a given analysis window, (e.g., the green line, representing a 1s window, at t=0 is the classification accuracy of a time window from −0.5 to 0.5 seconds). The time windows that achieved the highest accuracy for each window size are indicated by the shaded rectangles, and the individual confusion matrices are reported in **Fig. 5b**.

### 8. ROC analysis

Sensitivity is calculated as the ratio of true positive (TP) classifications to actual positive conditions, or the combination of TP and false negatives (FN).

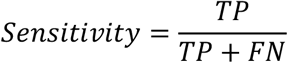

Specificity is calculated as the ratio of true negative (TN) classification to actual negative conditions, or the combination of TN with false positives (FP).

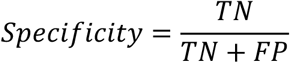

For non-binary classifications in which there are more than two possible outcomes we used a one-vs-rest method to convert into a binary classification. In short, the odor of interest is the positive odor, and all other odors are the combined negative (**Fig. 4d**).

The threshold for classification was varied by weighting the Euclidean distances calculated between testing and training templates (**Extended Data Fig. 4**). Distances between any testing template with the training template of the odor of interest were multiplied by the threshold weighting value. Then, the testing templates were assigned to the closest training template using the weighted distance values. Therefore, at a threshold weighting value of 1, this analysis is the same as using minimum Euclidean distance to assign the testing odor templates. Threshold weighting values greater than 1 reduce the chance for testing templates to be assigned to the odor of interest, and vice-versa for threshold weighting values less than 1. The sensitivity and specificity were calculated for a range of thresholds and plotted on a receiver operating characteristic (ROC) curve (**Fig. 6a,b**). The sensitivity and specificity across the range of thresholds of 100 random subsets of 10, 20, and 30 neurons from the recorded population of 32 PNs were averaged together to produce the ROC curves. This analysis was replicated for subsets of neurons ranging from 5 to 31. The area under the curve (AUC) for each subset of neurons is calculated and averaged. An exponential model fits the data with the y-intercept at 0.5 because no neural data leads to a completely random binary classifier, or an AUC of 0.5 (**Fig. 6c,d**).

Youden’s Index was calculated as a combination of sensitivity and specificity to find the best threshold weighting to use (**Fig. 6e,f**).

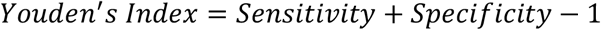

### 9. Photoionization detector

A commercial photoionization detector (PID) (Aurora Scientific, miniPID model 200B) with a 10.6 eV, RF-excited, lamp was used to check the VOC delivery profile of the olfactometer to the insect (**Extended Data Fig. 1c**). The built-in pump on the PID was set to the lowest setting (750 sccm) to pull air into the PID. After allowing the lamp to warm up for 30 minutes, the PID was calibrated to 0.1 μV using clean air with a gain of 5x. 80 sccm of headspace from the bacterial odor flasks were diluted in 720 sccm (10% dilution) for a total of 800 sccm of air being delivered to the PID inlet via the olfactometer. Odors were delivered using the same timing (four seconds) of odor stimulation as the electrophysiology experiments for five trials each with 1-minute interstimulus intervals. Voltage outputs were collected at 100Hz and saved as LVM files to a PC for subsequent analysis. Each trial was normalized by subtracting the baseline, calculated as the average voltage output during the first 30 seconds of each trial, from that trial. The odor was released 35-39 seconds after each trial started. The five trials were averaged together, and then a three-point moving smoothing filter was applied.

## Notes

### Competing Interest Statement

The authors have declared no competing interest.

