## Supplemental File for "Noninvasive detection of bacterial biofilms using an insect olfactory brain-based gas sensor"

**Extended Data Figure 1**

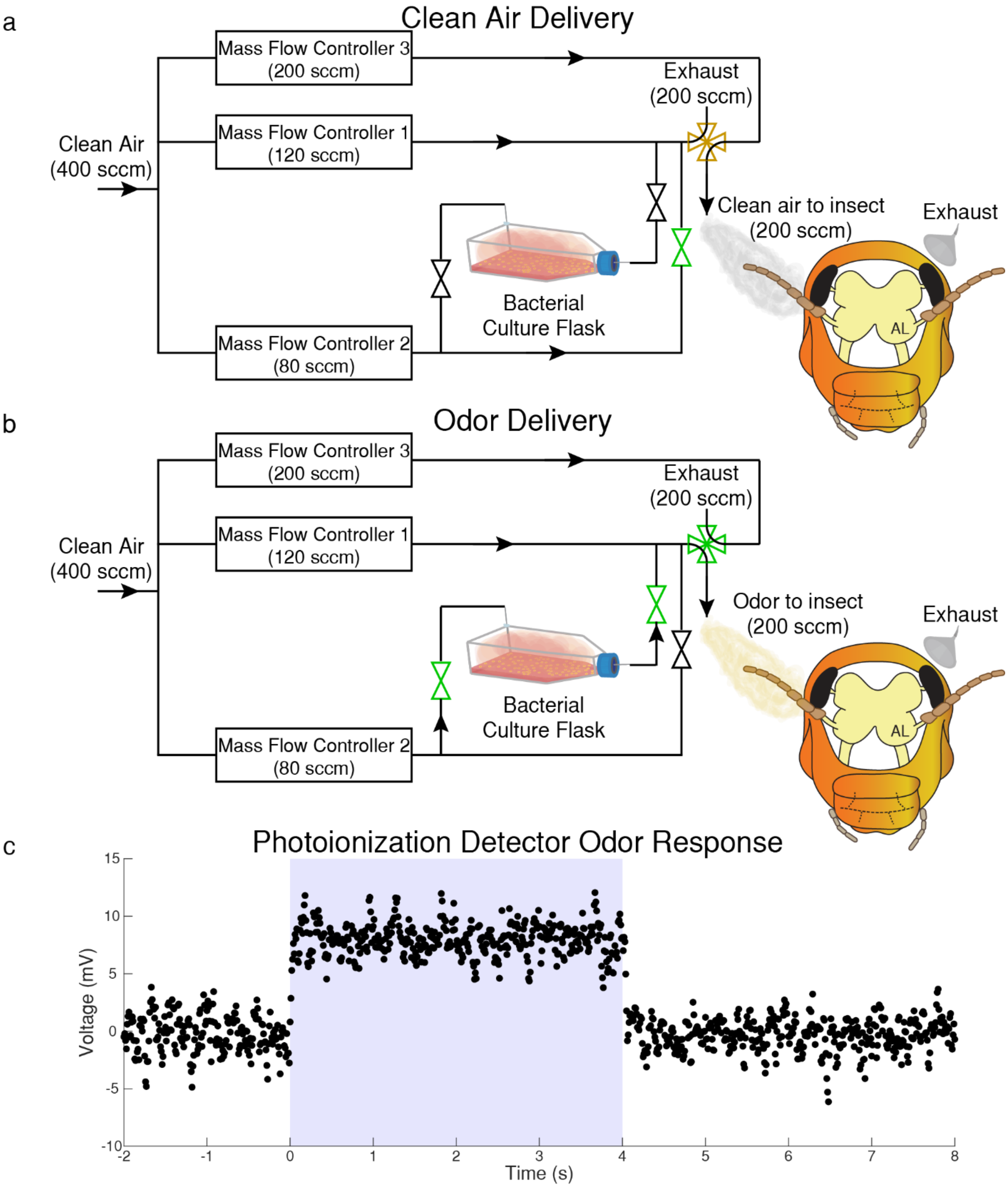

### Extended Data Figure 2

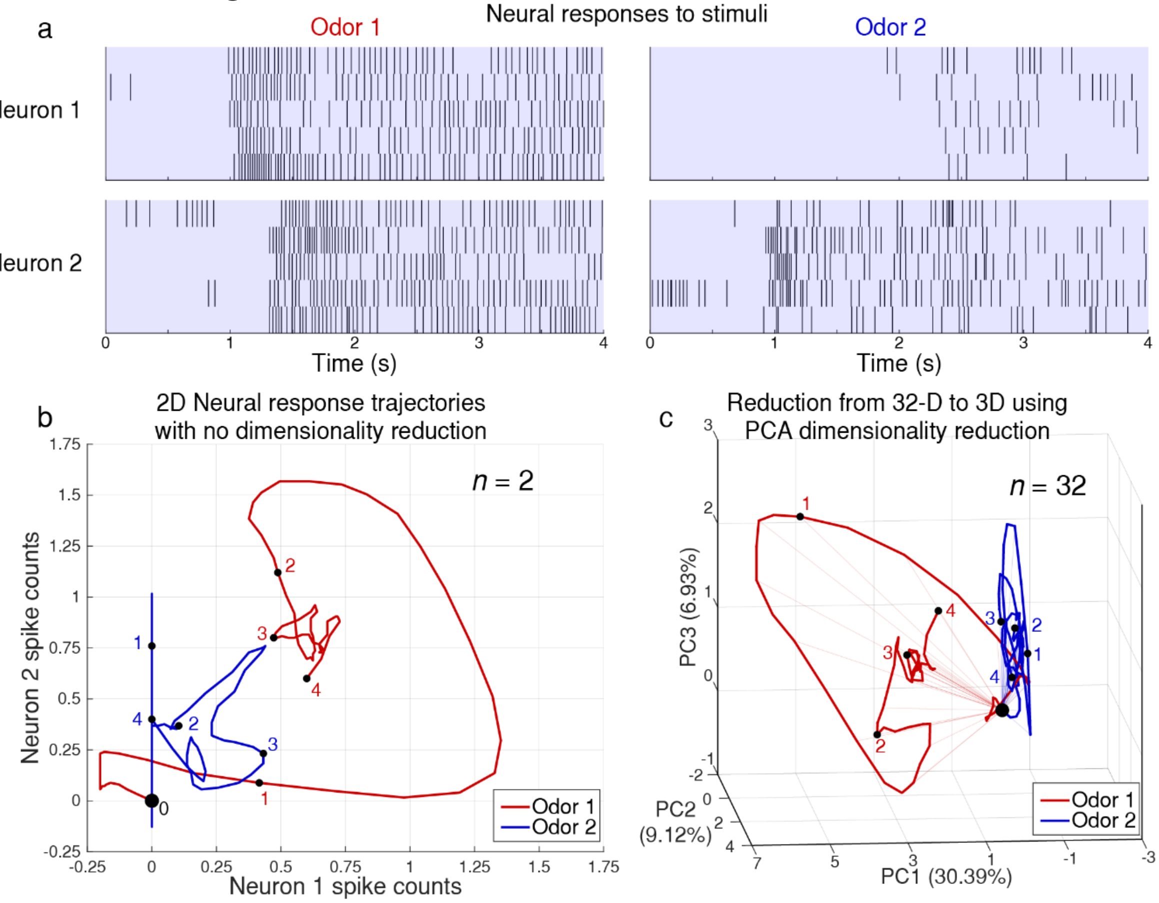

**Extended Data Figure 3**

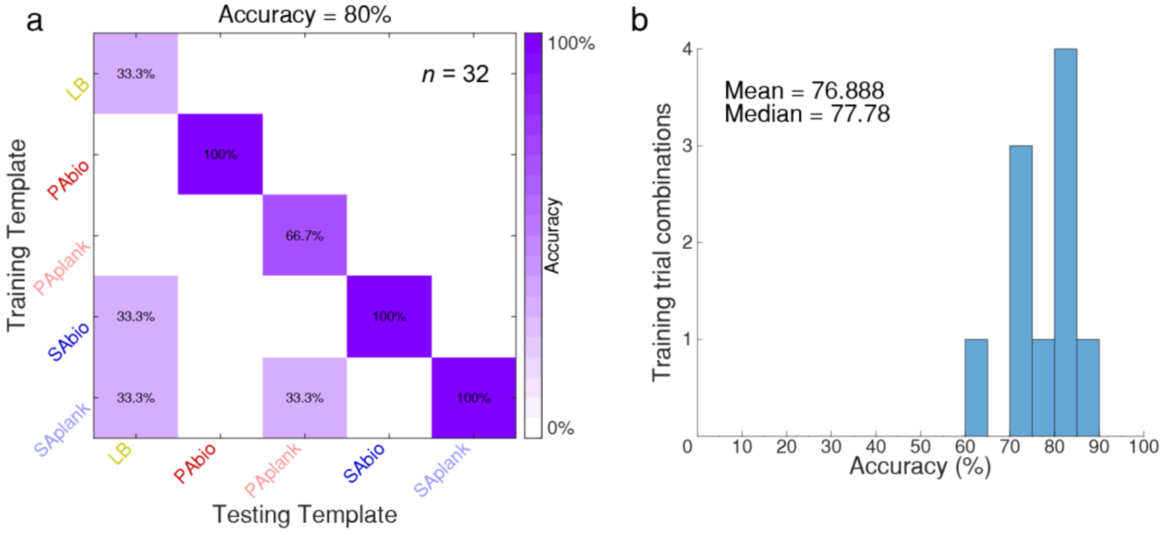

**Extended Data Figure 4**

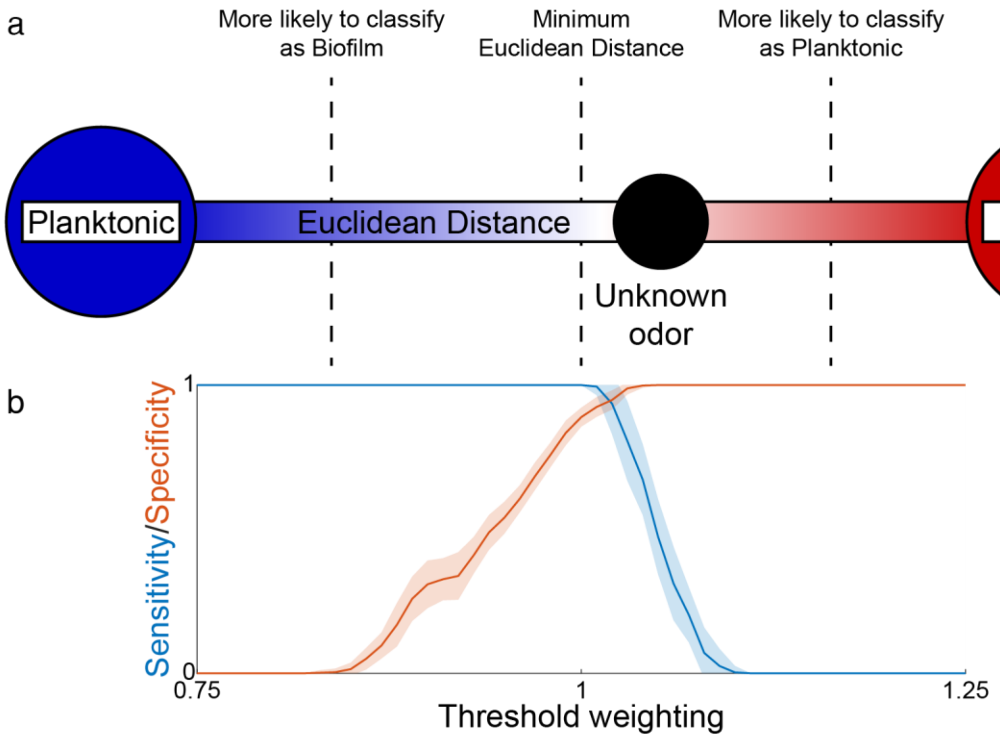

**Extended Data Figure 1: Air flow and odor delivery to the insect antennae is carefully controlled by the olfactometer. (a)** Prior to any odor delivery, the insect receives 200 standard cubic centimeters (sccm) of clean air via Mass Flow Controller 3 (MFC3). The final valve is a four-way valve that can switch back and forth between delivering the flow from MFC3 or the combined flows from MFC1 and MFC2. While clean air is being delivered to the insect antennae, the valves around the bacterial culture flask are closed to preserve the headspace containing VOCs, and the bypass valve connecting MFC1 and MFC2 is open. **(b)** During odor delivery to the insect, the final valve switches to deliver the combined airflows from MFC1 and MFC2. The valves around the bacterial culture flask are opened, and the bypass valve is closed to divert the 80 sccm of airflow from MFC2 through the bacterial culture flask. This 80 sccm of VOC laden-air mixes with the 120 sccm from MFC1, and this combined 200 sccm of air with bacterial odor in it is delivered to the insect antennae. This valve configuration is maintained for four seconds of the odor delivery before switching back to the clean air delivery setup. This setup maintains a constant airflow of 200 sccm to the insect, preventing possible mechanosensory neural responses from biasing the measured odor response. **(c)** Using a photoionization detector (PID), we confirmed the odor delivery profile using the LB control. The olfactometer odor delivery follows a square wave profile.

**Extended Data Figure 2: Neural trajectories visualized using PCA also include the temporal changes across time. (a)** The neural responses of two neurons to two different odors can be visualized using raster plots. Neuron 1 had a much stronger response (more spikes) to Odor 1 than to Odor 2. Also, Neuron 1's response to Odor 2 is delayed. Neuron 2 also had different responses between the two odors, with a slightly more delayed response and more consistent spiking to Odor 1. **(b)** We plotted the neural responses to each odor with each neuron along different axes and then connected the point in temporal order to reveal the neural trajectories to each odor. We saw clear differences in the curve of each neural response. **(c)** When using the neural responses from more neurons (n = 32), there are now 32 orthogonal axes, which we cannot visualize. We used PCA to reduce the dimensionality to three for visualization and then connected the points in temporal order to reveal the trajectories.

**Extended Data Figure 3: Using independent training and testing templates yields similar classification accuracies as LOTO analysis.** Instead of using LOTO (**Figs. 2d, 3d, 4b**), here we created training templates by averaging two random trials together, and then tested using the remaining three trials. **(a)** Using the WTA approach yielded 80% accuracy. **(b)** To check the robustness of these results, we then tested each of the possible combinations of two training trials and summarized the accuracies as a histogram. Nine out of the ten possible combinations (five trials, choose two trials) had accuracy between 70-90% with one combination resulting in a lower accuracy of 60%.

**Extended Data Figure 4: Varying the classification model threshold. (a)** The training templates for a planktonic and a biofilm odor can be visualized as opposite ends of a spectrum. In reality, these templates are in high high-dimensional neural encoding space. An unknown odor, or testing template, is also within this space. The distances between the unknown odor and each of the training templates can be calculated, and then the unknown odor assigned to whichever training template it is closest to. This would use the minimum Euclidean distance metric. However, not all classification tests want to equally weigh a positive (biofilm) and a negative (planktonic) classification. Therefore, the threshold, that was previously in the middle, can be shifted towards the biofilm, which would increase the chance of the unknown odor being classified as planktonic, or it could be shifted in the opposite direction. **(b)** As the threshold is weighed differently, the sensitivity and specificity change. At one extreme, for low thresholds, all unknown odors are assigned to the biofilm training template, which yields a sensitivity of 1, but a specificity of 0. For the opposite extreme, the sensitivity is 0, and the specificity is 1. However, as the threshold shifts, a region in the middle yields high sensitivity and specificity.
